# Machine Learning Based Modelling of Human and Insect Olfaction Screens Millions of compounds to Identify Pleasant Smelling Insect Repellents

**DOI:** 10.1101/2023.12.25.573309

**Authors:** Joel Kowalewski, Sean M. Boyle, Ryan Arvidson, Jadrian Ejercito, Anandasankar Ray

## Abstract

The rational discovery of behaviorally active odorants is impeded by limited information on how the olfactory system generates percept or valence for a volatile chemical. In previous studies, we showed that chemical informatics could be used to predict ligands for a large repertoire of odorant receptors in *Drosophila* (Boyle et al., 2013). However, it remained difficult to predict behavioral valence of volatiles since the activities of a large ensembles of odor receptors encode odor information, and little is known about the complex information processing circuitry. This is a systems-level challenge well-suited for Machine-learning approaches which we have used to model olfaction in two organisms with completely unrelated olfactory receptor proteins: humans (∼400 GPCRs) and insects (∼100 ion-channels). We use chemical structure-based Machine Learning models for prediction of valence in insects and for 146 human odor characters. Using these predictive models, we evaluate a vast chemical space of >10 million compounds *in silico.* Validations of human and insect behaviors yield very high success rates. The discovery of desirable fragrances for humans that are highly repulsive to insects offers a powerful integrated approach to discover new insect repellents.

## Introduction

Animals can detect volatile molecules in their environment using specialized protein receptors in their olfactory organs. Mammalian odorant receptors were discovered in 1991 and are G-protein coupled receptors (GPCRs) numbering >400 in humans (Buck L and Axel, 1991). Several years later starting in 1999 three families of distinct insect odorant receptors were discovered from the *Drosophila* genome which are ligand-gated ion channels and number ∼100 (Clyne et al., 1999, 2000). Over the subsequent 20-30 years, studies in mice and insects focusing on odor detection, odorant receptors, odorant receptor neurons, and higher-level circuitry, have revealed important principles. A single volatile is usually detected by a combination of receptors, and the activity from the neural circuit processing the information is extremely complex. Previously, we successfully developed a chemical informatics approach to discover ligands for a large number of odorant receptors in *Drosophila* (Boyle et al., 2013). However, the mechanisms connecting olfactory perception (e.g., rose, lemon, pleasant, unpleasant, etc.) to olfactory receptor activity are incompletely understood and it is therefore challenging to predict behavior from odorant receptor activity alone. There is also often a lack of correspondence between simple, 2D representations of odorants and their perceptual descriptions, and the molecular-cellular organization of the olfactory system lacks a simple discernible coding strategy that enables predictions, unlike auditory or visual processing (Buck L and Axel, 1991; Grabe & Sachse, 2018; P Mombaerts, 1999; Peter Mombaerts, 2001; Sachse & Beshel, 2016; Vassar et al., 1993).

Yet Machine Learning algorithms have shown promise in accurately predicting human odor perception (Keller et al., 2017; Kowalewski et al., 2021; Kowalewski & Ray, 2020b; Lee et al., 2023; Sanchez-Lengeling et al., 2019). Recently, we have shown that the volatile structure-to-percept prediction can be improved by using the activities of a small number of human odorant receptors and applying machine learning algorithms to identify highly predictive combinations (Kowalewski & Ray, 2020b). In insects, we have limited success modeling the behavioral valence of volatiles from olfactory receptor neuron activity in the fruit fly, *Drosophila melanogaster*. Since receptor activity depends on chemical structure, this suggests machine learning algorithms can potentially detect a consistent physicochemical basis for simple behavioral or complex perceptual responses (Kowalewski & Ray, 2020b; MacWilliam et al., 2018).

To test whether Machine learning based modeling can lead to identification of strong behaviorally active volatiles, we selected a challenge that has a very high potential for easing human suffering and yet had little innovation: pleasant smelling fragrances that are also repulsive to harmful insects that transmit diseases to humans or cause agricultural loss. Annually, mosquitos account for ∼500,000-700,000 deaths and a billion infections due to vector-borne illnesses like malaria, Dengue, Yellow Fever. Insects also destroy ∼30% of the global agricultural produce causing additional suffering. Insects are currently managed through widespread application of insecticides, but mutations are emerging that confer insecticide resistance and affect insect olfactory responses in both human disease vectors and plant pests (Deletre et al., 2019; Duneau et al., 2018).

The most widely used topical repellent for mosquitoes is DEET (N,N-Diethyl-m-toluamide), a chemical with undesirable chemical and cosmetic properties. DEET (N,N-Diethyl-m-toluamide) was discovered >75 years ago through empirical testing. It is not widely used outside the developed world, where vector borne diseases are present globally, and are also use restrictions due to its strong solvent properties, even dissolving plastics and nylons. Recent studies have also raised human health concerns, as the chemical interferes with inhibitory neurotransmission in mammals via GABA_A_ and Glycine receptors (Grant & Hall, 2019). Unfortunately, not enough is understood about the odorant receptors for aversion in mosquitoes as a result there has been limited innovation in the search for repellents even a quarter century after discovery of insect odorant receptors. The last repellent registered by the US EPA for commercial use was Picaridin, over 20 years ago.

Volatile compounds that have pleasant fragrances are desired by humans in cosmetics and are therefore excellent candidates for novel mosquito repellents. In order to identify fragrances that would be aversive to insects, we assumed machine learning (ML) could be used to identify structural features enriched among repellents and then detect these features within any arbitrary, large chemical space (Maldonado et al., 2006). We trained a support vector machine (SVM) on a comprehensive set of chemicals with known repellency from literature and identified predictive physicochemical features. In parallel, we adapted a collection of recently developed ML models to predict 146 different human odor percepts, helping us organize the candidate repellents by their potential fragrances (Kowalewski et al., 2021; Kowalewski & Ray, 2020b). We then validated the accuracy of behavioral predictions for insects and humans by randomly testing 50 predicted chemicals for repellency in mosquitoes and odor perception in humans. The validation success was >75% in each assay, repellent and perceptual, with ∼95% of the chemicals tested for repellency in round 1 showing aversion, which left ∼5% as neutral or slightly attractive to mosquitoes. Collectively, this suggests a strong physicochemical basis for odor valence and percept in insects and humans. In addition, we also built *in silico* toxicity models, filtering the compounds across multiple parameters for human use. A small number of the fragrances were tested on evolutionarily distinct insects, finding that they are repellent to mosquitoes and the fruit fly, *Drosophila*.

Tens of millions of dollars and several years may be required for identification and adequate human-safety analysis for registration of new repellent chemicals by the EPA (Gupta & Bhattacharjee, 2006). Machine learning approaches for modeling toxicity as well as other physical and cosmetic attributes are steadily advancing, suggesting it is possible to accelerate this search and significantly reduce costs. Accordingly, we included additional computational filters to select predicted repellents such as predicted fragrant odor perceptual qualities. Because fragrant molecules that are also insect repellents may be rare to find, we applied this approach to a large library (>10 million) of purchasable chemicals from the ZINC 15 database. These efforts represent a dramatic expansion of the insect repellent chemical space and provides a theory-based high-throughput method to drive rational discovery of novel insect repellents.

## Results

### Machine learning can learn to predict aversive valence in insects from chemical structure

To predict insect behavior, we developed a novel chemical informatics approach using Machine Learning (ML). We assembled a training set of volatile chemicals known in the literature to have a negative behavioral valence including eucalyptol, linalool, geraniol, citronella, alpha-thujone, beta-thujone (Kline et al., 2003; Klocke et al., 1987; Syed & Leal, 2008), DEET, picaridin, 34 N-acyl piperidines that are structurally related to picaridin (Katritzky et al., 2008) and structurally diverse odorants from fruits and food sources that are not expected to elicit aversion (Carey et al., 2010; Hallem & Carlson, 2006). We generated energy-minimized 3D structures for each and calculated 3,242 physicochemical features for 201 training chemicals. From it we identified 18 features that were highly correlated with aversive valence (correlation of 0.912). We then fit a support vector machine (SVM) with these features to learn to predict the repellency of test chemicals (Figure 1).

**Figure 1.**
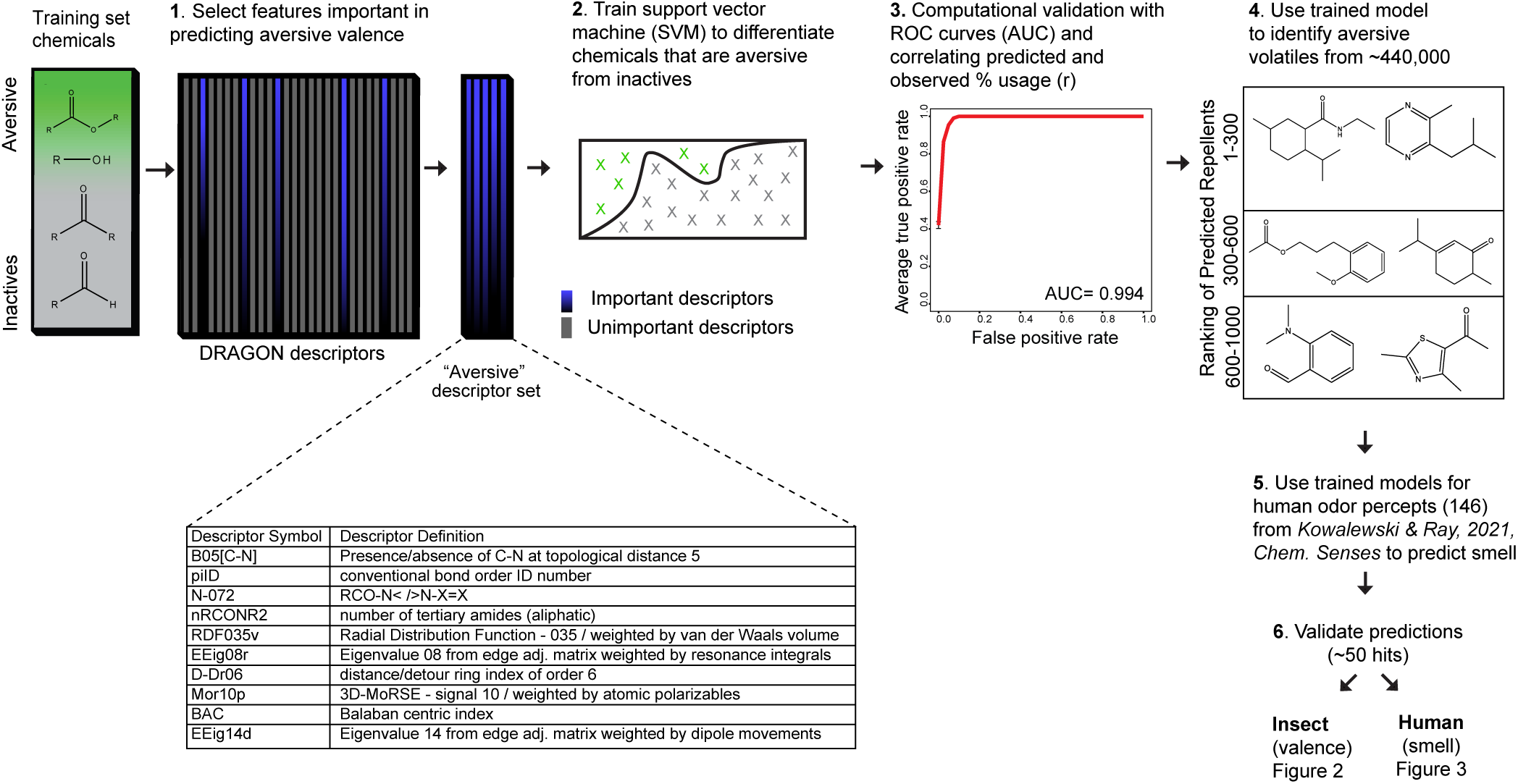
A chemical informatics pipeline to screen for pleasant smelling insect repellents. **A**, Overview of the cheminformatics pipeline used to identify novel DEET-like ligands from a larger chemical space. Summary table of top 10 physicochemical features that optimally corelate with repellency. Receiver-operating-characteristic curve (ROC), which indicates the success in predicting test repellents. The performance is the average over 100 80/20 splits, where 80% of the data is for training and 20% for testing. The mean area under the curve (AUC) is provided. Representative structures from the top 150 predicted repellent compounds.

The trained model successfully predicted the valence of the test chemicals (avg AUC = 0.994), indicating that it could distinguish volatiles with aversive valence from non-repellents (Figure 1). Evaluation of the predictive chemical features indicated six membered rings, carbon-nitrogen distances, tertiary amides, and the positioning of oxygen atoms in the molecule are associated with repellency (Figure 1).

We subsequently used this repellency model to evaluate a >440,000 small molecules assembled from databases which includes ∼3,000 chemicals from natural sources like plants, insects, vertebrates, and compounds that are in food, beverages, and cosmetics. The top 500 predicted compounds (top ∼0.1% of hits) represent a structurally diverse group of chemicals, with some showing similarities to volatiles in the training set (Figure 1). Several naturally occurring compounds were also in the list of precited repellents. Since the ML model successfully classified known repellents versus non-repellents, we next experimentally tested the candidate repellents.

### Machine learning can prioritize candidate repellents by modeling their odor qualities to humans

Importantly, repellents must meet many physical and cosmetic demands for human use. However, evaluating the scent characteristics of candidate repellents with human subjects is a laborious, expensive, and time-consuming task. In order to create a method to bypass this bottleneck, we adapted our recent perceptual modeling approach involving the prediction of 146 odor perceptual descriptors from chemical structure; example perceptual descriptors include “fruity other than citrus”, “lemon”, “orange”, “cinnamon”, “perfumery” and unpleasant ones such as “sickening”, “rancid”, “animal”, and “dirty linen” to name a few (Kowalewski et al., 2021; Kowalewski & Ray, 2020b). The models were trained with data from prior human behavior studies in which participants evaluated ∼150 odorants and mixtures according to the 146 descriptors. The development of the methods and validation of the models were rigorously evaluated in the previous publications (Kowalewski et al., 2021; Kowalewski & Ray, 2020b). Using these models, we predicted odor descriptors for the candidate insect repellents that were discovered using the model described in Figure 1. The compounds had a wide variety of predicted odor characteristics covering both pleasant and unpleasant fragrance profiles. Together with the insect valence predictions, this gives us the foundations for selecting desirable fragrances that can protect humans and plants from pests.

### Validation of computational behavioral predictions of human smell and insect aversion

To validate the behavioral predictions of scent or repellency, on humans and insects, respectively, we selected 44 compounds based on their availability and affordability. First, we compared them to reports from human testing of smell properties. While odor qualities remain poorly characterized, 37 of the 44 tested had odor and taste annotation in a sensorial databases at the The Good Scents Company. We could then rigorously evaluate the accuracy of the trained perceptual models by ROC analysis. The perceptual models predicted the observed descriptors for most volatiles with a high success rate (Avg. AUC = 0.77) (Figure 2A,B). Interestingly, many chemicals in the predicted repellents have pleasant odor characteristics, including highly desirable ones such as fruity, floral or perfume-like, such as seen in the word cloud of enriched predicted characters for the predicted insect repellents (Figure 2C).

**Figure 2.**
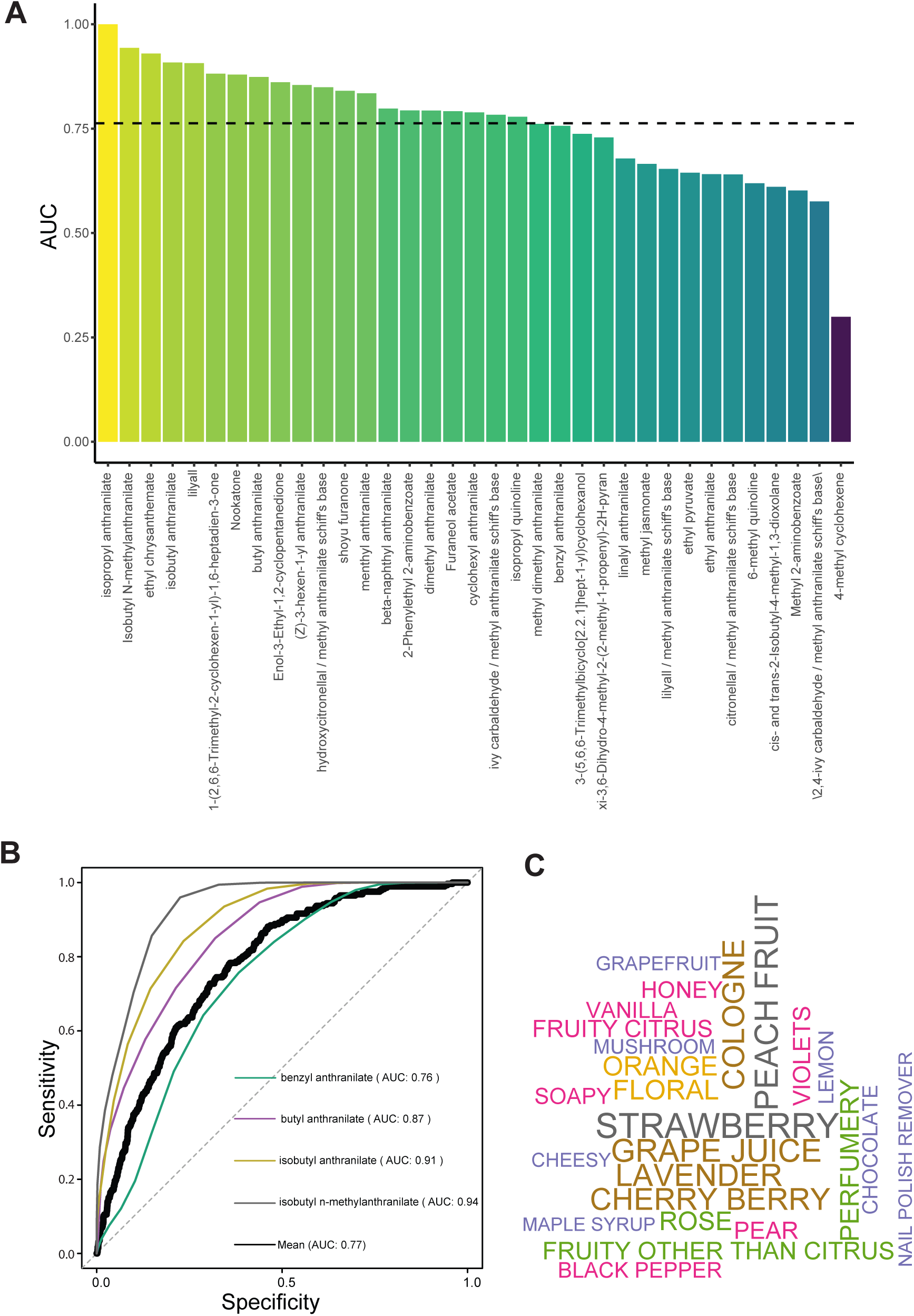
Models successfully predict odor perceptual qualities of repellents in humans. **A,** Validation of models predicting the odor perceptual qualities (146 perceptual descriptors) of a set of compounds that were screened for repellency in Figure 3A. The performance metric for the validation is the area under the ROC curve, which assesses the rate of correctly predicted perceptual descriptors relative the rate of incorrectly predicted descriptors. Dashed line is the average Area Under the Curve (AUC). **B,** Sample ROC curves for select compounds as well as the ROC curve across all compounds (e.g., shown as separate curves in the ROC plot as well as those not shown but whose identities appear in Figure 2A); the respective AUC values are reported inside the plot. **C,** The top 25 perceptual descriptors for the predicted repellents that underwent experimental validation are displayed as a word cloud. The font size of the descriptor is scaled relative to frequency.

We next tested the compounds against *Aedes aegypti* mosquitoes. We developed a high throughput assay utilizing the innate attraction of the female mosquitoes to a heat block approximating human skin temperature (37 °C) covered with a filter paper (Figure 3A). Each of the test compounds were individually evaluated by applying on the filter paper (3% in acetone) and counting the numbers of mosquitoes landing on the surface with solvent control before and after applying the compound. A repellency index indicated that most of the predicted compounds (∼90%) showed >50% repellency to female *A. aegypti* mosquitoes (Figure 3B). These results suggest that the machine learning (ML) algorithm was capable of predicting a chemical’s perceptual qualities for humans and its valence for insects. This set of chemicals represents a very large increase in new insect repellents, comprising diverse chemical structures and smell characteristics that are desirable for human use. In a separate study we had tested a few of these repellents using a human arm-in-glove behavioral assay and observed strong repellency (Boyle et al., 2016), suggesting that the newly discovered compounds are likely to exhibit similar effects.

**Figure 3.**
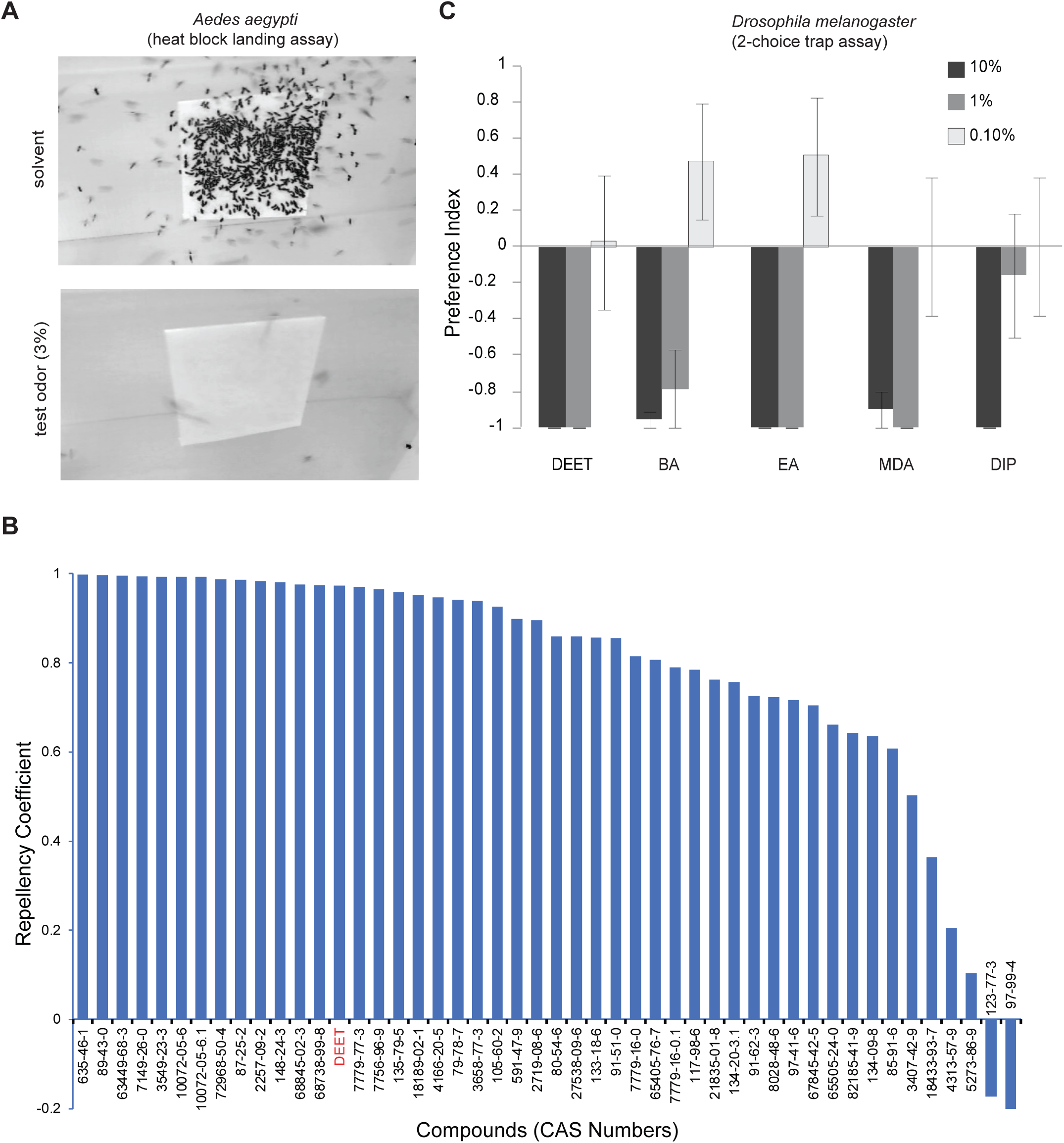
Experimental validation of predicted insect repellents show a high success rate several of which are pleasant smelling. **A,** Representative still photograph of landing assay of female *Aedes aegypti* mosquitoes on filter paper covered heat block over 5 mins (overlay of all frames), and **B,** mean Repellency Coefficient for each of the indicated compounds (by CAS numbers) tested at ∼0.2mg/cm^2^, N=2-4 trials for each, ∼20 mosquitoes/trial. For DEET, N=23 trials. CAS numbers ending in a “.1” are compounds tested at a second lower amount of ∼0.06mg/cm^2^. **C,** Mean Preference index of *Drosophila* adults to repellents at three different concentrations dissolved in acetone in a Two-choice trap assay baited with 10% Apple Cider Vinegar measured after 48 hrs. N = 7-10 trials each treatment at 48 hrs, 10 flies/trial, error bars = s.e.m.

Since we do not know the molecular receptor targets for these diverse repellents, and different insect species have both highly diverse and conserved olfactory receptors, we cannot tell *a priori* whether the repellency is narrowly effective in mosquitoes or conserved across diverse insects. To determine broad-spectrum aversive valence among insects, we tested another distantly related dipteran, *Drosophila melanogaster*, which is >250 MY apart evolutionarily. All of the predicted compounds displayed concentration-dependent aversive valence to *Drosophila* in a trap-based 2-choice assay (Reeder et al., 2009; Syed et al., 2011). Most of the tested compounds showed strong repellency (Figure 3C). The simplest interpretation is that the machine learning algorithm was able to identify chemical features that predict negative valence across multiple species of dipteran insects.

### Repellents show calcium mobilization in mammalian cells

An alternative approach to identifying insect repellents would be to computationally model a known cellular or molecular activity relevant to repellency. Interestingly, one of the aversive compounds used in the repellency training set is the commercial repellent DEET, had previously been seen to cause calcium mobilization in human HEK293 cells at high concentrations (Dennis et al., 2018). In order to further evaluate if such Ca^2+^ mobilization is a common feature for repellency, we performed additional experiments.

When HEK293 cells were exposed to DEET, a dose-dependent calcium transient was observed, with an EC50 of 56.6 mM (∼1% vol/vol), below the concentrations used on skin applied formulations (7-98% vol/vol) of which up to 60% is absorbed into the skin (Schoenig et al., 1996; Selim et al., 1995). (Figure 4A). Additional cell lines from different organisms were tested, including an insect S2 cell line, and all of them were sensitive to DEET at similar EC50s (Figure 4A). The DEET-induced calcium transients were similar in magnitude to the positive control, a 100 µM ATP induced signal (e.g., GPCR mediated) (Figure 4B). To assess the source of DEET induced calcium flux, cells were depleted of intercellular calcium with treatment with thapsigargin (a SERCA antagonist) 10 minutes before the assay. These calcium-depleted cells did not show increased Ca transients when treated with DEET. This indicates that DEET is likely inducing release from intercellular calcium stores, in a dose-dependent manner (Figure 4C). Interestingly, two additional commercially used repellents, picaridin and diethyl benzamide (a structural analog of DEET), also evoked a strong intracellular Ca^2+^ signal (Figure 4D). Because these three compounds were part of the training data to build our machine learning insect repellent models, we tested whether the newly identified aversive fragrances also showed similar activity. We tested a sampling of the verified aversive fragrances, and a few traditionally known repellents, at two concentrations (5% and 0.5%) in the HEK293 cell line and found that there were indeed some that showed Ca^2+^ release (Figure 4E). However, there appeared to be no correlation between the repellency index and the Ca^2+^ activation of the fragrant repellents (R^2^=0.0093), suggesting that this conserved effect is unlikely to contribute to aversive behavior.

**Figure 4.**
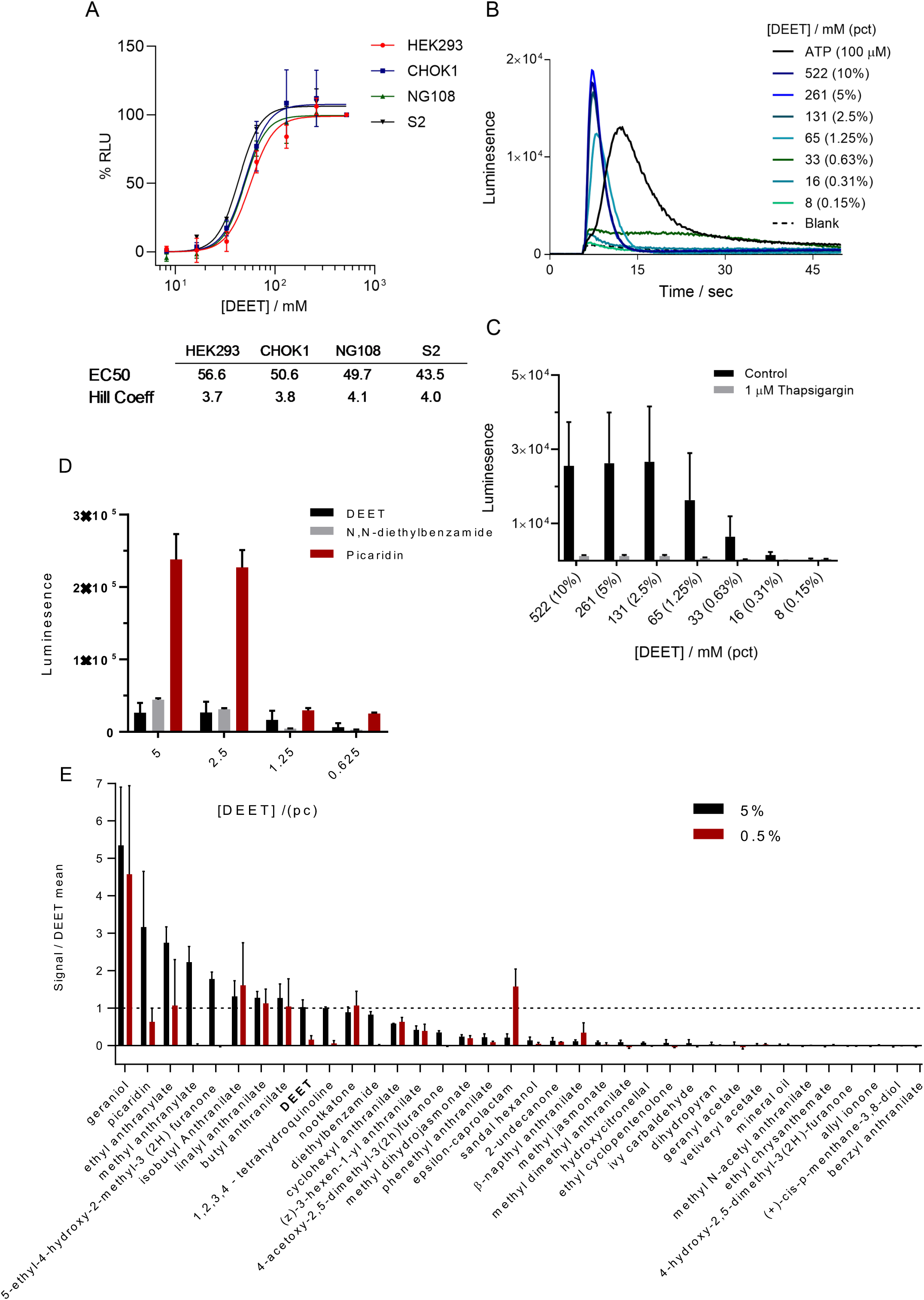
Some, but not all insect repellents increase Ca^2+^ mobilization in a human cell line. DEET induces a non-specific calcium response *in vitro*. **A.** Dose response curves for three mammalian cell lines (HEK293. CHOK1, NG108) and one insect cell line (S2). EC50 and Hill coefficients for each cell line are given below. **B.** Representative DEET induced calcium kinetics in HEK293 cells. **C.** HEK293 cells treated with thapsigargin 10 min before assay to deplete intracellular calcium stores no longer respond to DEET, suggesting DEET response is dependent on intracellular calcium. Error bars are standard deviation. **D. F,** Luminescence of the Ca^2+^ indicator at two different concentrations for predicted and previously known repellent chemicals, scaled relative to DEET.

Nevertheless, the data provided an opportunity to develop a better understanding of the physicochemical basis of the Ca^2+^ mobilization seen *in vitro* and we trained machine learning models to predict the calcium activity (Figure S1A). Upon analyses, the top chemical features for predicting the calcium activity were different from the features selected by the model to predict repellency. This too suggest that while there is a structural basis for basis for calcium activity which does not overlap with shared features for detection as a repellent. The features fell into 3 broad categories including 2D autocorrelation, RDF, and 3D MoRSE features. Autocorrelations are distance-based metrics, with the 2D autocorrelation referring to the number of bonds between a pair of atoms in the molecule, while radial distribution function (RDF) features encode dependencies between atoms that give rise to molecules’ characteristic infrared spectra (Hemmer et al., 1999). The 3D MoRSE features are based on theoretical electron diffraction patterns and are therefore sensitive to differences in 3D geometries (Devinyak et al., 2014). In previous work, these features have been shown to be predictive of ligand bonding and biological activity (Beglari et al., 2020; Cheng et al., 2010; González et al., 2006).

Evaluation of models trained with these features on test data showed a high correlation between the predicted and observed Ca^2+^ activities (Figure S1). Using another approach, where we classified the chemicals as broadly falling into one of two categories (high versus low activity), the predictions were also accurate, further emphasizing that the Ca^2+^ signal does indeed have a strong physicochemical basis. Collectively, this suggests the unique 3D shape, rather than generic, possibly indiscriminate information, underlies the observed Ca^2+^ activity, supporting our contention that the activity may be a persistent off target effect among certain chemical repellents and should therefore be further investigated. While the physiological consequences of this effect have not been studied, we incorporated this activity as a parameter into our machine learning pipeline, allowing us to further triage repellents for future use by low estimated Ca^2+^ activity, in addition to other desirable characteristics for human or animal use.

### Mining massive commercially available chemical spaces for novel insect repellents

The size and diversity of the chemical library we initially screened (∼0.45M) is in the range of high-throughput wet lab screening that pesticide discovery companies can undertake. The Machine learning approach allows us to screen far more without the limitations of cost of synthesis and screening time. To demonstrate the massive throughput capacity of this approach, we increased our search space to >10 million commercially available chemicals (from the ZINC 15 database), canvasing far more structural diversity than the ∼450,000 chemicals we previously screened. We built a new training set including the chemical repellents we experimentally verified, more than doubling the number of the actives while adding greater structural diversity. The top features fell mostly into 2D and 3D autocorrelations and 3D MoRSE categories, suggesting that certain types of features became increasingly important following the addition of more repellent activities. Notably, some features changed from this vastly improved training set; that is, moving away from simple features that account for amides, or more generally, chemical bonds including nitrogen (N). This possibly reflects an initial bias toward DEET, an amide, that was resolved when including more complex feature selection methods and increasing the size and diversity of the training set.

There was a strong correlation between the predicted and observed repellency when validating these updated models on test data (R = 0.73). Then, by applying the filters that we progressively incorporated into the pipeline, and described throughout (e.g., insect valence, odor perceptual qualities, and intracellular calcium release), we identified a set of candidates still numbering in the hundreds. We next predicted the acute oral mammalian toxicity (Rat) using a computational model from our earlier work (Kowalewski & Ray, 2020a) and eliminated all compounds falling into EPA Tox categories I and II. The top predictions contain many aminobenzoates comprised of benzene, amino, and ester groups. Some chemicals notably differ in that they resemble amide repellents like diethyltoluamide (DEET) (Figure 5A). The top hits were predicted to have a variety of different human odor characters, many of which are perceptually pleasant or are useful in agriculture (Figure 5B). This approach accelerates the identification of novel, fragrance-based, broad spectrum insect repellents.

**Figure 5.**
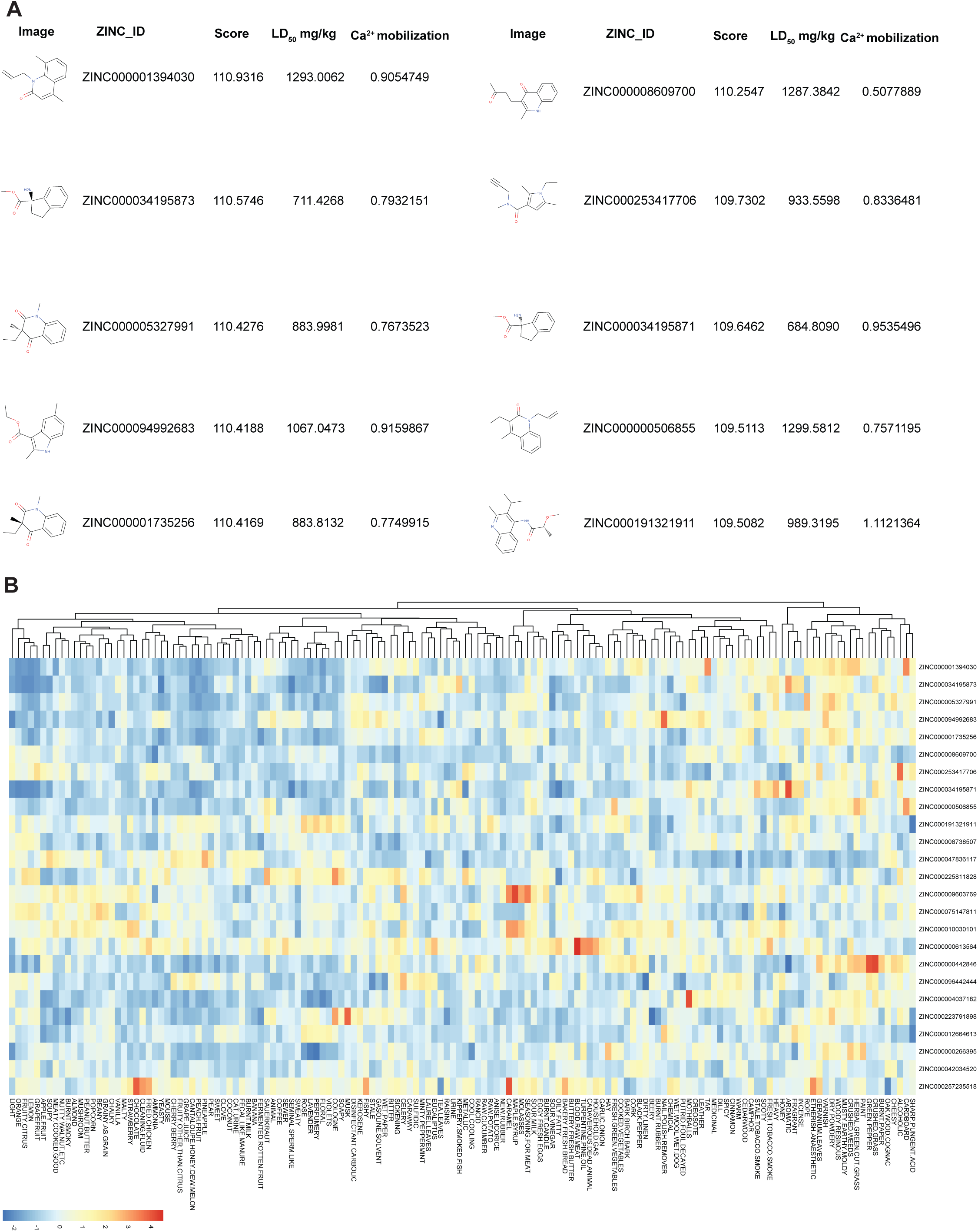
Screen of 10+ million chemicals predicts several insect repellents with different smells. **A,** Tabulated predicted % repellency (relative to DEET) from a library of 10+ million chemicals, filtered to the top values. The best matching known repellent is displayed alongside the Euclidean distance, the predicted LD50 and Ca^2+^ mobilization. **B**, heatmap organizing top predictions according to estimated perceptual qualities.

## Discussion

Here, we have developed a comprehensive machine learning pipeline to screen small or large chemical spaces for suitable alternatives and verified numerous predicted chemicals for repellency in various behavioral assays and across species, indicating the effectiveness of the pipeline. The rapid identification of fragrances that have excellent behavioral activity, pleasant smell and safety greatly improves the likelihood of regulatory approval and potential for use.

Interestingly, we identified a strong intracellular Ca^2+^ signal in response to DEET and picaridin as well as some anthranilates. While the mechanism remains unclear, it has been previously observed in cultured cell lines after DEET exposure (Dennis et al., 2018; Swale et al., 2014). One possibility proposed in the literature is that the elevated intracellular Ca^2+^ is due to activation of octopamine receptors. Octopamine plays a modulatory role, with targets including OctαR and OctαβR in insects and TAAR1 in mammals (Kleinau et al., 2011; Swale et al., 2014), and while this could explain the many complex physiological responses that have reported in the DEET literature (Briassoulis, 2001; Clem et al., 1993; Corbel et al., 2009; Tavares et al., 2019), an unknown but conserved receptor subtype is still possible. Additional research will be needed to determine whether this phenomenon is important for off-target mammalian toxicity. In either case, the model we developed to predict the Ca^2+^ mobilization may help isolate a set of key physicochemical features, enabling identification of compounds that have this effect. However, since the calcium activity does not explain the repellency, additional electrophysiological and genetic experiments are needed to further clarify the mechanism of action.

Although many repellents have been studied, DEET remains the most widely used commercially available chemical repellent. Yet research continues to emerge about off-target toxicities in humans who are repeatedly exposed to DEET; specifically, accounts suggest adverse skin reactions and activities in human or broadly mammalian nervous systems (Briassoulis, 2001; Grant & Hall, 2019; Osimitz et al., 2010). This study demonstrates the successful development and application of a machine learning pipeline that accelerates research into insect repellency and its physicochemical basis. The methods and data presented provide a high-throughput and rational way to identify additional novel repellent chemicals. Here, we emphasized screening candidate chemicals that are most likely to fit multiple requirements (smell and repellency) rather than simply repellency. As these requirements steadily increase, it is obvious that massive, unexplored chemical spaces offer the most promising leads. By predicting and filtering 10+ million purchasable chemicals according to their desirability for human use, we further illustrate an essential role for computational approaches (e.g., machine learning) in future repellency studies, accelerating the discovery of safe, ecologically friendly, and effective repellent chemicals.

## Material and Methods

### Behavior testing

#### Repellency Assay in Drosophila

Repellency was tested using a *Drosophila melanogaster* Canton S strain (RRID:BDSC_1), backcrossed >5 generations to w1118 (RRID:BDSC_3605) strain in 2-choice trap assay. All chemicals were purchased at highest purity from Sigma Aldrich (RRID:SCR_008988), TCI, Alfa Aesar, Penta, Parachem, Bedoukian, and Molbase. A chamber was built from DRAM vials, with four air holes made in the plastic lid and the bottom of chamber is coated with 5 ml of 1% agarose to maintain moisture. Traps are made from 1.5 ml Natural Microcentrifuge tubes with bottom 3 mm cut to form an opening and a 1000 ul blue graduated pipette tip catalogue number 1111-2023. To create a plastic funnel, a pipette tip was cut on both ends; cut to 1.0 cm from tip end to expand the opening of the funnel and then cut on the other end of plastic to create a final height of 2.5 cm for funnel. A BioRad PCR 0.2 ml Tube cap was cut from the strip and was inserted into the lid of microcentrifuge tube. Test odor (35 ul) was placed in the PCR tube cap and lure (90 ul) was placed in outer ring between the PCR cap and the microcentrifuge cap. In the dark, 4–7-day old, starved flies (10 males, 10 females) were released per trap and followed for 2 days. Counts were taken manually, and the Preference Index (PI) was computed using the formula below.

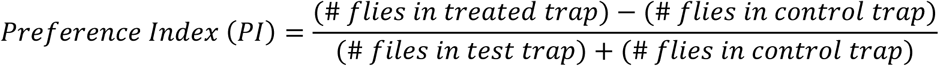

#### Repellency Assay in Mosquitoes

A non-contact heat attraction assay was performed by simultaneously offering mated female *Aedes aegypti* mosquitoes that were ∼4-10 days old, a test chemical on a filter paper coupled with an attractive heat source. Heat for the assay was provided by a Hot hands® Hand warmers HH2 (Heat Max, Dalton, GA) which was activated by shaking in gloved hands. Temperature (∼37°C) was verified using IR thermometers. The hand warmer was fitted snugly into a 100 mm x 15 mm petri dish base and covered with 15 cm x 15 cm piece of polyester net secured round the petri dish by a pair or 8-inch plastic cable ties (Gardner Bender, Milwaukee, WI). The petri dish with hand warmer was covered with a larger 150 mm X 15 mm petri dish with a 10 cm x 7.5 cm rectangular window cut out. This netted window aligned with the hand warmers exposing a heated surface.

A polyester net (measuring 9 cm x 8 cm) was treated with 500 µl of either acetone (solvent) or 3% (v/v or w/v) test chemical and allowed to air dry for 30 minutes to evaporate solvent. The test compound (∼0.2mg/cm^2^) treated net was secured by stretching between a frame made using two 10 cm x 7.5 cm flexible magnetic frames with 7.5 cm x 6.2 cm cut out windows. This dose is 88% lower than the amount recommended in the WHO, WHOPES “Guidelines for Efficacy Testing of Mosquito Repellents for Human Skin” recommended dose (∼1.67mg/cm^2^) actives. Three more flexible magnets were added on top as spacers from the heat source. The net-magnet assembly was placed directly over the 10 cm x 7.5 cm cut out openings of the larger 150 mm X 15 mm petri dishes holding the heat pads. The stimulus was placed on the top net of the frames. Temperature was monitored by a laser Infra-red thermometer and assay started at 37°C. A puff of 5% CO_2_ was blown into the cage to behaviorally activate the mosquitoes and initiate the assay. A webcam timelapse is used to record mosquitoes landing on the net for automated counting offline using ImageJ (RRID:SCR_003070). A custom Macro is used in ImageJ (RRID:SCR_003070) to count the mosquitoes in the selected area (the net) across the timepoints. The counts from each frame are automatically tabulated.

#### Macro

run(“8-bit”); setAutoThreshold(“Default”);

//run(“Threshold…”);

setThreshold(0, 80);

run(“Analyze Particles…”, “size=2-70 show=[Overlay Outlines] clear summarize stack”);

dir = getDirectory(“image”);

name = getTitle;

index = lastIndexOf(name, “.”);

if (index!=-1) name = substring(name, 0, index);

name = name + “.csv”;

selectWindow(“Summary of “+getTitle);

saveAs(“Measurements”, dir+name);

Since this is a 1-choice assay, for each cage and compound, a solvent alone control is done before and after the test material, with at least 20 minutes between the trials. Any significant changes in behavior between the control solvent should indicate if mosquito behavior is acceptable.

*Repellency coefficient* = (mean cumulative number of mosquitoes on solvent treated control netting – mean cumulative number of mosquitoes on treated netting)/ mean cumulative number mosquitoes attracted to solvent treated control netting. Cumulative number of mosquitoes on the window of treatment are counted for at time points 2,3,4,5 min. Each compound has 2-23 trials, ∼40 mosquitoes per trial.

#### In vitro assay of Ca^2+^ mobilization

All mammalian cell lines were maintained at 37 C, 5% CO2, and 99 percent relative humidity. Insect Schneider 2 (S2) Cells (RRID: CVCL_Z232) were maintained at 26 C and 99% relative humidity. HEK293T (ATCC, RRID:CVCL_0063), CHOk1 (RRID: CVCL_E8IB) and NG108-15 (RRID: CVCL_0464) were grown in DMEM media (Gibco), supplemented with 10% FBS (Gibco), and antibiotic-antimycotic supplement (Gibco). S2 cells (RRID: CVCL_Z232) were cultured in Shields and Sang M3 Insect Medium (Sigma), supplemented with Insect Medium Supplement (Sigma). Cells were transfected with the luminescent calcium reporter aequorin 24 hours before experiment with X-tremeGENE 9 transfection reagent, in the case of HEK293T (and CHOK1, and X-tremeGENE HP, in the case of NG108 and S2 cells. Transfected cells were harvested with enzyme-free cell dissociation buffer (Gibco), centrifuged, and suspended in assay media (DMEM with 0.1% BSA for mammalian cells, unsupplemented SFX-100 insect medium for S2). The aequorin cofactor coelenterazine-F (Nanolight) was added (5 µM final concentration) to the cell suspension, in the dark, with gentle stirring for 2 to 3 hours before assay. For water-insoluble compounds like DEET an emulsion was prepared by adding a percentage of DEET indicated, vol/vol to assay media, vortexed, and bath-sonicated for at least 15 min, but up to one hour. Cells (100 µl, at approx. 50,000 cells) were injected into a 96 well plate containing 100 DEET µl emulsion, and resulting luminescence recorded on a BMG Labtech Lumistar Omega plate reading luminometer. Thapsigargin was added to the cell suspension were indicated to a final concentration of 1 µM, 10 minutes before assay.

### Chemical Informatics

#### Machine learning predictions of insect repellency

Physicochemical features for figures 1-2 were selected using Sequential Forward Selection (SFS) (Haddad et al., 2008) from 3,224 using Dragon software (Talete). This algorithm is an iterative search over the chemical feature pool that seeks to maximize the correlation between chemical feature values and, here, the % repellency. These chemical features were subsequently used to fit a support vector machine (SVM) with a radial basis function kernel (RBF). The implementation of the support vector machine algorithm is from the e1071 R package. We performed 20 independent 5-fold cross-validations followed by a receiver operating characteristics (ROC) analysis to analyze the performance of the repellency prediction method. The trained SVM then ranked both a library obtained from eMolecules (http://www.emolecules.com) (>400,000) and natural odor libraries (Supplementary Material) (>3,000) by the repellency score estimates and the set of optimally predictive chemical features. The EPI Suite (http://www.epa.gov/oppt/exposure/pubs/episuite.htm) was used to calculate predicted LogP and Vapor Pressure values.

For Figures 3, 5, S1-S2, we updated the methods and algorithms to further optimize the pipeline. Physicochemical features were selected using AlvaDesc software, which offers > 5000 chemical features. Optimal chemical features were then selected using cross-validation and recursive feature elimination algorithm (CV-RFE). The RFE CV algorithm involves iterative partitioning of the data and the selection of feature subsets (n_1_, n_2_, …, n_i+1_; where “n” is the size of the feature subset and “i” is an arbitrary indexer). Given a data partition and n features, an arbitrary ML algorithm is fit, importance is assigned to the feature set and the model of n features predicts a test partition, that is, data excluded from training, differentiating the CV-RFE from RFE alone. In this study, the ML algorithm was a support vector machine (SVM) or a random forest. Over 100s of iterations the algorithm converges on the optimal (n) number of features to maximize predictive success. Additionally, for every round the contribution of each chemical feature is assessed, giving rise to an aggregate rank; namely, the frequency at which a feature has been identified as important to the predictive success. Again, distinguishing the standard RFE algorithm from CV-based adaptation. As for computing importance, the random forest algorithm internally tests the contribution of the features. It accomplished this by fitting multiple decision trees on bootstrap samples of the data. Consequently, a percentage of data has been excluded and serves as a test set. By predicting the test set with and without permuting or shuffling the values of a given feature, the algorithm can then assign an importance according to the % increase in error that is observed after permuting the feature.

The support vector machine (SVM) algorithm does not natively perform a comparable bootstrapping procedure as random forest and therefore does not generate a test set to assess feature importance. In this case, an external metric must be supplied to assign the importance. Depending on the type of fit, classification or regression, this is a pseudo R^2^, based on non-linear regression, or the AUC for the feature and the outcome that is being predicted (e.g., % repellency). For additional details, refer to the documentation provided with caret package in R.

#### Machine learning predictions of Ca^2+^ mobilization

Due to the smaller number of observations (datapoints or chemicals) we opted for an ensemble method that could take advantage of more of the available data. Random forest fits multiple decision trees on bootstrap samples of the data, thereby reducing the tendency to overfit to one or few chemicals in the dataset. As compared to the SVM, it has two simple parameters (mtry and ntrees) and the tuning grid (or number combinations of these parameters that need to be tested to optimize performance tends to be small). Accordingly, it is an ideal approach when little is known about the activity being predicted.

To further reduce overfitting, we identified a small set of physicochemical features on different subsets of the data and then randomly sampled from this pool. By doing so, the features are slightly underfit, which is the better approach, again, when less is known about the activity being predicted.

Additionally, we used two different methods and metrics to independently verify that physicochemical features were indeed predictive of the activity. First, a regression-based approached predicting the raw assay activities. Second, a classification-based approach where the top ∼40% of the activity distribution was converted into an “Active” label, assessing predictive success by ROC analysis. Finally, we fit classification models that were otherwise identical but trained on shuffled labels. This ensured that the chance level-performances significantly differed.

### Machine learning predictions of odor perceptual qualities

We used methods that were developed previously and extensively validated for prediction of human odor characters from chemical structures and are described there (Kowalewski et al., 2021; Kowalewski & Ray, 2020b). Briefly, perceptual training data was from a study published in a reference book: ATLAS of Odor Character Profiles (Dravnieks, 1985). In the study, panelists supplied ratings for ∼150 odorants and mixtures across 146 possible descriptors; the ratings are reported as the % usage. The % usage refers to the portion of panelists that rated an odorant (1-5) using a particular descriptor (1-146), indicating its relevance to the odorant. It is on the scale 0-100, with the maximum of 100% suggesting every participant found the descriptor to be relevant. The top perceptual descriptors were selected by thresholding the predictions based on the mean value in the training data. For each descriptor in the training data, thresholds were computed as 2x the mean. These thresholds were then applied to the predicted repellents to generate a set of exemplar descriptors. Subsequently, these exemplars were converted into a descriptor frequency table, where the frequency is the total number of occurrences of a descriptor. The data and word cloud plot were generated using the R programming language, with additional functionality provided by the Wordcloud package in R.

## RESOURCE AVAILABILITY

### Lead Contact

Anandasankar Ray

### Materials Availability

No new materials were generated in this study.

### Data and Code Availability

Data used in the analyses is publicly available from the references cited, and the data generated in this manuscript are supplied within the figures. Any additional data associated with this manuscript is also available in other formats on request from the corresponding author and lead contact.

The code developed for this study is available from the corresponding author for non-commercial use upon request. Use of the code may be subject to a standard Material Transfer Agreement (MTA) or specialized licensing agreement through University of California Riverside’s Technology Transfer Office, as patent applications related to this software are currently pending.

## ACKNOWLEDGMENTS

This work was partially funded by University of California Riverside internal HATCH funding and partially funded by NIDCD and NIAID. The content is solely the responsibility of the authors and does not represent the official views of the funding agencies. The funders had no role in study design, data collection and analysis, decision to publish, or preparation of the manuscript.

## AUTHOR CONTRIBUTIONS

J.K., S.M.B., R.A. and A.R. conceived the study. J.K. and S.M.B. planned and executed the computational analysis. R.A. planned and executed the cell line experiments and J.A. performed the insect behavior experiments. J.K. and A.R. wrote the manuscript with contributions from others.

## FIGURE LEGENDS

**Supplemental Figure S1.**

**A**, Overview of the approach to generate and validate the machine learning models. Here, a model-averaged prediction is made, where each model has access to different physicochemical features and is trained on different combinations of training set chemicals. **B,** The R^2^ values over the 100 train/test splits, averaged into 10 bins. **C,** The average classification success is reported over 100 train/test splits, assessed by the area under the ROC curve. Active labels (positive cases) were assigned as chemical scoring in the top ∼40% of Ca^2+^ values. This is compared to shuffling the labels before training, reported as “Shuffle.”

