## Supplementary figures and images for "Machine Learning Based Modelling of Human and Insect Olfaction Screens Millions of compounds to Identify Pleasant Smelling Insect Repellents"

### Supplemental Figure S1

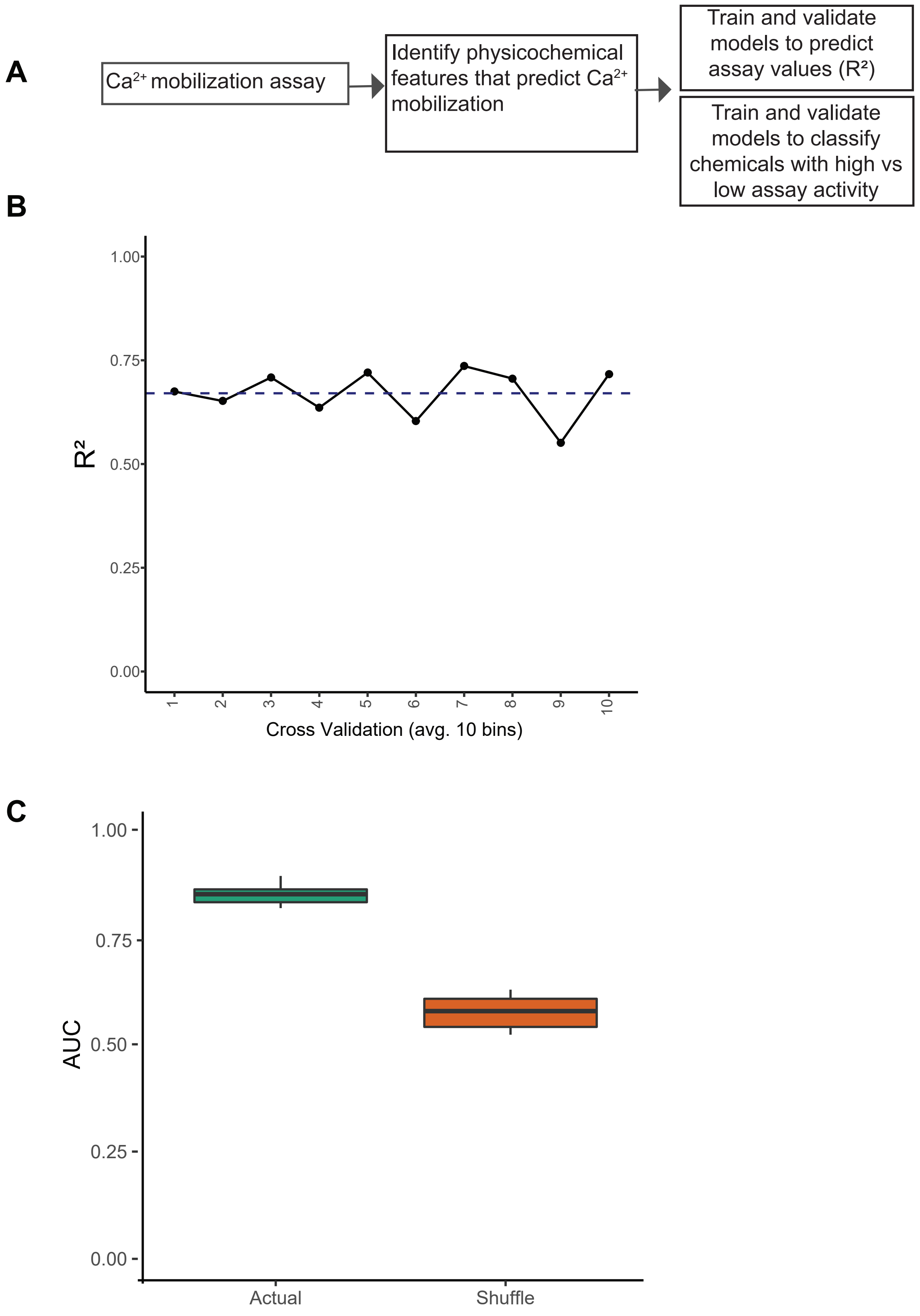
